# Machine learning based gut microbiota pattern and response to fiber as a diagnostic tool for chronic inflammatory diseases

**DOI:** 10.1101/2023.03.27.534466

**Authors:** Miad Boodaghidizaji, Thaisa Jungles, Tingting Chen, Bin Zhang, Alan Landay, Ali Keshavarzian, Bruce Hamaker, Arezoo Ardekani

**Affiliations:** School of Mechanical Engineering, Purdue University, 585 Purdue Mall, West Lafayette, IN 47907, USA; Department of Agricultural and Biological Engineering, Purdue University, West Lafayette, IN 47907, USA; Departments of Internal Medicine, Anatomy and cell biology,, and Molecular Biophysics and Physiology, Center for Integrated Microbiome and Chronobiology Research, Rush University Medical Center, Chicago, USA; State Key Laboratory of Food Science & Technology, Nanchang University, Nanchang, China; School of Food Science and Engineering, South China University of Technology, Guangzhou 510640, China

**Keywords:** Microbiome data, Machine learning, Ulcerative colitis, Crohn’s disease, Fiber treatment

## Abstract

Gut microbiota has been implicated in the pathogenesis of multiple gastrointestinal (GI) and systemic metabolic and inflammatory disorders where disrupted gut microbiota composition and function (dysbiosis) has been found in multiple studies. Thus, human microbiome data has a potential to be a great source of information for the diagnosis and disease characteristics (phenotypes, disease course, therapeutic response) of diseases with dysbiotic microbiota community. However, multiple attempts to leverage gut microbiota taxonomic data for diagnostic and disease characterization have failed due to significant inter-individual variability of microbiota community and overlap of disrupted microbiota communities among multiple diseases. One potential approach is to look at the microbiota community pattern and response to microbiota modifiers like dietary fiber in different disease states. This approach is now feasible by availability of machine learning that is able to identify hidden patterns in the human microbiome and predict diseases. Accordingly, the aim of our study was to test the hypothesis that application of machine learning algorithms can distinguish stool microbiota pattern and microbiota response to fiber between diseases where overlapping dysbiotic microbiota have been previously reported. Here, we have applied machine learning algorithms to distinguish between Parkinson’s disease, Crohn’s disease (CD), ulcerative colitis (UC), human immune deficiency virus (HIV), and healthy control (HC) subjects in the presence and absence of fiber treatments. We have shown that machine learning algorithms can classify diseases with accuracy as high as 95%. Furthermore, machine learning methods applied to the microbiome data to predict UC vs CD led to prediction accuracy as high as 90%.

## Background

Microbial communities play invaluable functional roles in supporting human health, including immune, metabolic, brain, and behavioral traits [1]. The human gastrointestinal tract harbors the largest population of micro-organism community, known as gut microbiota. Gut microbiota typically is composed of a variety of micro-organisms, including bacteria, archaea, and eukarya [2]. The gut microbiota actively participates in the human metabolism, contributing to the synthesis of vitamins and other nutrients, regulating immune functions, promoting gut barrier integrity, among others [3]. Not surprisingly, several disease-related states have been linked with a disbalance (i.e., dysbiosis) of the gut microbial community [4]Whether or not gut microbiota dysbiosis is the cause, the consequence, or both of these in different diseases states, it contains invaluable information to aid in the diagnostics of diseases. Given the high dimensionality of microbiome data, there might exist hidden patterns or information that are hard to detect with the existing analytical methods.

Notably, however, is the variation of the gut microbiota of different individuals which not only differ according to disease states, but also genetics and several environmental factors, including diet and dietary fiber intake which change between and within individuals and populational groups [5]. Such individual gut microbiota variations prevent disease diagnostics based on clustering through commonly used beta diversity ordination plots that often show overlap across conditions with no clear group separations between healthy and disease states [6]–[8].

Recently, Machine learning (ML) methods have opened new doorways for the exploration of gut microbiota data in ways that were once impossible. Supervised and unsupervised machine learning methods have been used for classification, regression, clustering, and non-negative matrix factorization of the data [9]. ML models have been successfully implemented to distinguish healthy subjects from ones with gastrointestinal (GI) diseases, such as inflammatory bowel diseases (IBD) [10]. Furthermore, ML methods have shown promising results in predicting the diseases that do not directly affect the GI or occur outside the GI tract, such as cardiovascular diseases [11].

Although there is a myriad of studies that have utilized ML for gut microbiota disease predictions, most of these studies have focused on distinguishing healthy from non-healthy subjects or predicting diseases that fall within similar categories of diseases, such as predicting GI disease, including IBD and esophagus diseases [12], [13]. As a result, it is vital to see if the predictive capability of the algorithms is limited to certain classes of diseases or if this capability can be extended to simultaneously, for example, predict multiple GI and systemic diseases where dysbiosis with overlapping microbiota community characteristics is present and what conclusions can be made of that as previous studies have reported an overlapping dysbiotic microbiota, for example, for Parkinson’s disease and IBD [14]. Additionally, whether it is possible to distinguish between the diseases after the subjects are given treatments, particularly fiber treatments, has not been investigated. As a result, in the current study, we aim to apply machine learning techniques to make predictions using the data that correspond to five different conditions, including Parkinson’s disease (PD), Crohn’s disease (CD), ulcerative colitis (UC), human immune deficiency virus (HIV), and healthy control (HC) subjects in the absence and presence of fiber treatments.

## Materials and Methods

### Materials

Fecal samples from 10 healthy individuals (HC), 10 Parkinson’s disease patients (PD), 7 inactive Crohn’s disease patients (CD), 7 inactive ulcerative colitis patients (UC), and 2 HIV patients (HIV) were received from Rush University Medical Center. The samples were frozen at -80 °C and shipped overnight with dry ice. Five fermentable soluble dietary fibers commonly present on diets were used for the study. Fructooligosaccharides from sugar cane (Nutraflora, Ingredion, USA), barley beta-glucan (P -BGBM, Megazyme, Bray, Ireland), apple pectin (AF 710, Herbstreith&Fox Inc., Germany), sorghum arabinoxylan (extracted as previously described, Rumpagaporn et al., 2015 [15]), and a mixture of the four (fructooligosaccharides, beta-glucan, pectin and arabinoxylan, 25% each). We selected these disease groups because prior studies have shown that they all are associated with dysbiotic microbiota characterized by relative abundance of “pro-inflammatory” bacteria like bacteria belonging to protobacter phyla and decreased relative abundance of “anti-inflammatory” bacteria including short chain fatty acids (SCFA) producing bacteria. Thus, there was an overlap in changes in their microbiota community that resulted in difficulty to use microbiota data as a diagnostic and/or disease prediction tool.

### *In vitro* fecal fermentation

The fecal samples were thawed in the anaerobic chamber 30 min prior to the in vitro fecal fermentation experiment. The fermentation procedure used was similar as mentioned before (Kaur et al., 2011 [16]) except that all the procedures were conducted in an anaerobic chamber instead of using CO_2_ flushing. Dietary fiber (1%) and 5% feces were added to 5 mL of PBS buffer in an anaerobic tube. The tubes were sealed and incubated at 37 °C for 12 h. After 12 h fermentation, the fermenta were collected and centrifuged at 14,000 g for 5 min and DNA was extracted from the pellet using FastDNA™ SPIN Kit for feces (116570200, MP Biomedicals, USA). Informed consent was obtained from all donors and experiments were approved by the ethical committee Purdue University (IRB 1509016451). Further, all participants signed the Rush University Medical Center (RUMC) Institutional Review Board approved informed consent forms (ORA#: 07100403; 12020204; 07092603; L04092807)

### DNA sequencing and data preprocessing

The DNA obtained before and after *in vitro* fecal fermentation of stools was analyzed by 16s ribosomal RNA (rRNA) sequencing, which was performed by the DNA Services Facility at University of Illinois at Chicago. Briefly, the V3-V4 region of the extracted DNA was amplified using the 341f/806r primer set. The amplicon was detected by agarose gel electrophoresis. A second polymerase chain reaction (PCR) analysis was performed on the common sequences with primers containing Illumina adapters, a sample-specific barcode (10 bases), and linker sequences (called common sequences) at the ends of the forward and reverse primers. After 2 stages of PCR, the amplicons were sequenced using an Illumina MiSeq sequencer. The obtained raw sequences, which contain forward R1 and reverse R2, were merged. Chimeras were removed using USEARCH algorithm and sequences were then merged into one FASTA file and subject to UPARSE pipeline for operational taxonomic unit (OTU) clustering [17]. The taxonomic information for each OTU was determined using a ribosomal database project (RDP) classifier [18]. For the preprocessing, the OTU based data was normalized with respect to the cumulative count of all the microbiota for each subject before feeding into ML models. Then, the genus level data was used for ML analysis.

### Machine learning modeling

For all the classification problems in this study, we used four different ML algorithms, including the random forest (RF), support vector machine (SVM), artificial neural networks (ANN), and convolutional neural network (CNN). These methods have been successfully implemented to solve many problems that involve genomic datasets. [12]. All of these methods can be utilized for binary and multi-label classification purposes, such as diagnosing healthy-vs-non-healthy and healthy-vs-Parkinson’s disease-vs-colitis, respectively. To implement SVM and RF, we used scikit-learn classifiers [19] in python, where the one-vs-one scheme is used for multi-label classification. To implement ANN and CNN, the Multi-Layer Perceptron (MLP) classifier of scikit-learn [19] and PyTorch [20] were used, respectively. Additionally, to prevent unbiased comparison, the same preprocessed data was fed to each ML model. To find the best model parameters and compare different machine learning models, we used 5-fold cross-validation technique, where we used the average values of the predictions of all the 5 folds to report classification metrics. Furthermore, we reported all the well-known classification metrics, including the macro and micro F1, recall, precision, and accuracy when comparing the models. The micro and macro-metrics were averaged over instances and categories, respectively [21].

In the SVM method, a hyperplane is used to separate data with a large amount of margin, and eventually, for the given kernel function, a hyperplane is found to classify the data into multiple groups. In general, SVMs work well when a clear margin of separation exists, and they can efficiently learn complex classification functions and employ powerful regularization principles to avoid data over-fitting [12]. SVM has shown promising capabilities to classify healthy and non-healthy subjects, for instance, in the case of lung cancer [22] or obesity [23]. In this study, we used the SVM algorithm with the non-linear radial basis functions as the kernel function.

RF utilizes an ensemble average of multiple decision trees, with each one working based on the bootstrapping method. In the case of classification, RF takes the majority vote of the trees for classification. Ensemble learning paves the way for learning complex and simple functions. Additionally, RF automatically assigns importance to each feature, which is used for selecting the important part of the spectrum. The major advantage of the RF method is the capability to handle datasets with a large number of predictor variables [24]. Furthermore, in general, RF does not require a comprehensive grid search for hyper-parameter optimization and the default parameters lead to acceptable accuracy [12]. RF has been applied to the gut microbiota for disease classification such as bipolar disorders [25], coronary artery disease [26] and major depressive disorder [27]. In the current study, we set the number of trees and maximum depth to 100 and 10 and the Gini impurity was used to calculate the quality of split.

Inspired by biological neural networks, deep learning methods such as ANN and CNN consist of multiple hidden layers and numerous neurons, which have the capability to be applied to a wide range of problems. Unlike most ML methods, neural networks have a built-in feature selection mechanism that assigns importance to each feature by multiplication by weights and applying activation functions. Further, in the case of CNN, applying convolutional layers enables the detection of spatial and dependencies in the input signals. Both DNN and ANN have been applied to different classification problems involving gut microbiota, such as obesity [28], inflammatory bowel disease [29], and Parkinson’s disease [30] detection. Different varieties of CNNs with different levels of data preprocessing, such as Met2Img, and Metal ML, which involve data augmentation and feature extraction, have been formed and applied to gut microbiota data [31]. Here, after some preprocessing and arranging the data in OTU forms, we directly fed the data into ANN and CNN. The architecture of CNN and ANN used in this study are shown in *Figure 1*. The cross-entropy loss function was used with both CNN and ANN, where the output is the probability for each one of the classes.

**Figure 1:**
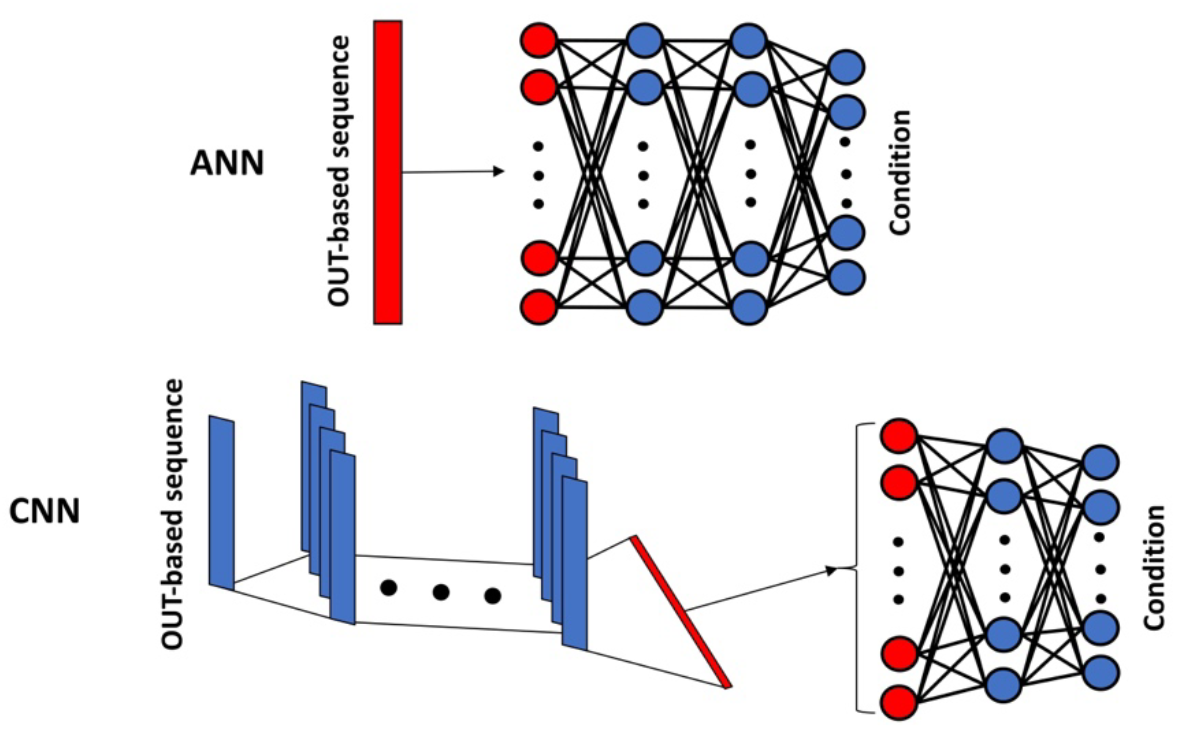
The schematic view of the ANN and CNN architecture used to predict the patient’s conditions. The number of neurons and layers shown here are for illustration purposes and do not reflect the actual values

## Results and discussions

We used two different datasets: the first dataset, where no treatments are given, and the second dataset, which comprises cases with and without fiber treatment. Throughout the manuscript, we refer to the first and second dataset as “baseline” and “fiber” datasets, respectively. For both cases, we changed the data size and monitored how the classification accuracy varies as a function of data size, as shown in Figure *2*. For both datasets, the prediction accuracy increased with the data size and reached values above 95%. For the classification purpose, we used the maximum data size, where 138 and 1092 data points were available for the baseline and fiber datasets, respectively. Additionally, since the values of micro metrics, including micro F1, recall, precision, and accuracy, are all the same in this study, we only reported accuracy as representative of all the micro metrics.

**Figure 2:**
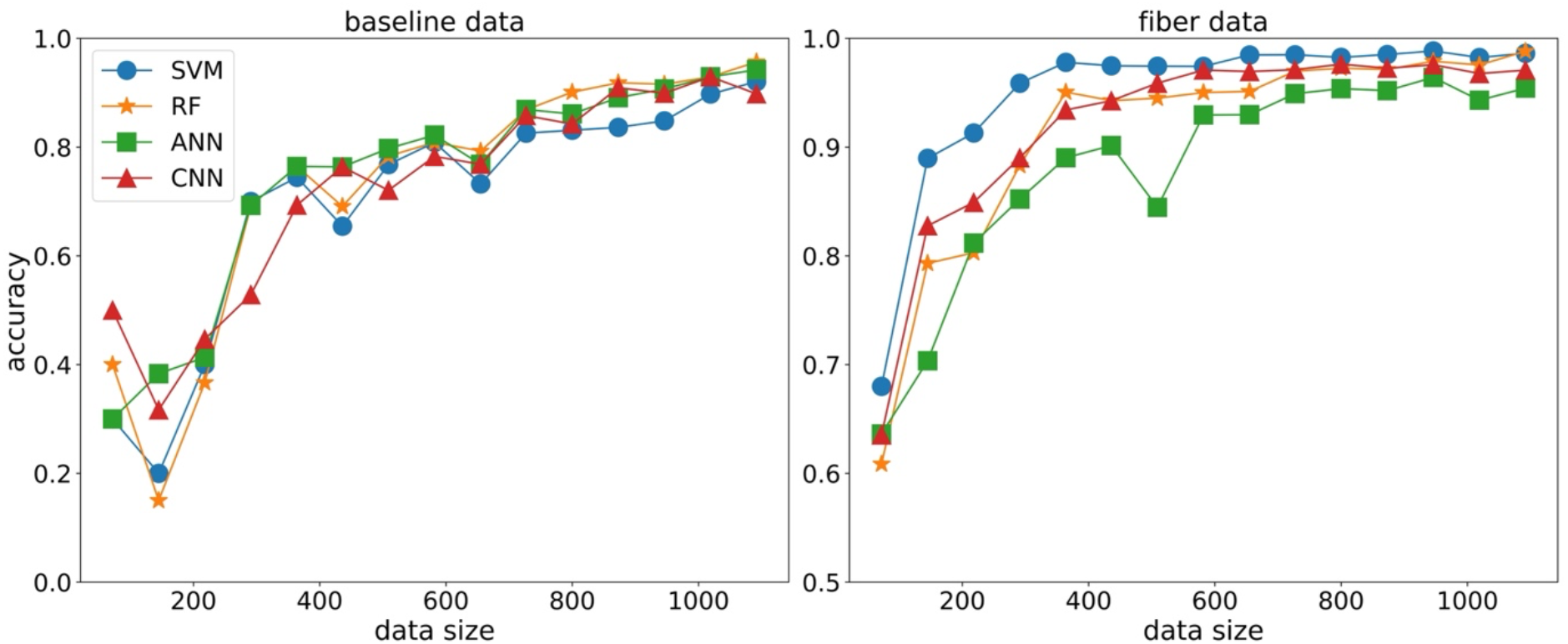
Prediction accuracy as a function of data size for the baseline and fiber data

### Prediction of the diseases based on the baseline data

Here, we present the results for three types of classification: 1) the five conditions, 2) the HC vs non-healthy (NH) 3), and the UC vs CD. Table *1* demonstrates the prediction performances for the baseline data. The results suggest that SVM provides the highest values for all the classification metrics. However, the difference between the algorithms is not significant. The heatmap for the classification of the five conditions for one of the folds in the 5-fold cross-validation is shown in Figure *3*. We note that algorithms have difficulty identifying UC vs CD and HIV vs PD diseases. The data points corresponding to HIV cases are relatively low in the dataset compared to other conditions, which contributes to the inability of the algorithms to detect HIV cases. For the UC-CD cases, we observed that classification metrics are not able to perfectly distinguish between the two, which is in line with what previous machine learning studies suggest. Indeed, among the different datasets and ML techniques used in the literature to diagnose UC vs CD, such as using RNA sequencing data [32] or endoscopic images [33], none has been able to perform as a perfect classifier. Additionally, we note that in the case of HC and NH, all the accuracies are relatively high for all methods, suggesting the strong capability of ML methods to distinguish between HC and NH. As the values in Table *1* demonstrate, the algorithms behave as almost perfect classifiers, where the macro and micro metrics lead to results as high as 99%. In other words, ML algorithms can perfectly identify the trend that distinguishes HC and NH cases, which is further reflected in AUC values, as shown in Figure *4*.

**Table 1:** Classification performances using the baseline data

| Task | Classifier | Macro precision | Macro Recall | Macro F1 | Micro | Accuracy |
| --- | --- | --- | --- | --- | --- | --- |
| Five conditions | RF | $0.890 \pm 0.113$ | $0.888 \pm 0.103$ | $0.885 \pm 0.110$ | $0.957 \pm 0.035$ | $0.957 \pm 0.035$ |
| | SVM | $0.975 \pm 0.022$ | $0.947 \pm 0.051$ | $0.952 \pm 0.045$ | $0.971 \pm 0.027$ | $0.971 \pm 0.027$ |
| | ANN | $0.953 \pm 0.034$ | $0.937 \pm 0.036$ | $0.940 \pm 0.033$ | $0.949 \pm 0.029$ | $0.949 \pm 0.029$ |
| | CNN | $0.958 \pm 0.035$ | $0.942 \pm 0.039$ | $0.946 \pm 0.035$ | $0.957 \pm 0.027$ | $0.957 \pm 0.027$ |
| HC-NH | RF | $0.995 \pm 0.011$ | $0.989 \pm 0.022$ | $0.991 \pm 0.017$ | $0.993 \pm 0.015$ | $0.993 \pm 0.015$ |
| | SVM | $1.000 \pm 0.000$ | $1.000 \pm 0.000$ | $1.000 \pm 0.000$ | $1.000 \pm 0.000$ | $1.000 \pm 0.000$ |
| | ANN | $1.000 \pm 0.000$ | $1.000 \pm 0.000$ | $1.000 \pm 0.000$ | $1.000 \pm 0.000$ | $1.000 \pm 0.000$ |
| | CNN | $1.000 \pm 0.000$ | $1.000 \pm 0.000$ | $1.000 \pm 0.000$ | $1.000 \pm 0.000$ | $1.000 \pm 0.000$ |
| UC-CD | RF | $0.892 \pm 0.047$ | $0.863 \pm 0.045$ | $0.865 \pm 0.051$ | $0.869 \pm 0.049$ | $0.869 \pm 0.049$ |
| | SVM | $0.913 \pm 0.049$ | $0.887 \pm 0.077$ | $0.883 \pm 0.077$ | $0.887 \pm 0.072$ | $0.887 \pm 0.072$ |
| | ANN | $0.874 \pm 0.101$ | $0.867 \pm 0.102$ | $0.866 \pm 0.102$ | $0.867 \pm 0.101$ | $0.867 \pm 0.101$ |
| | CNN | $0.860 \pm 0.087$ | $0.847 \pm 0.081$ | $0.847 \pm 0.083$ | $0.849 \pm 0.082$ | $0.849 \pm 0.082$ |

**Figure 3:**
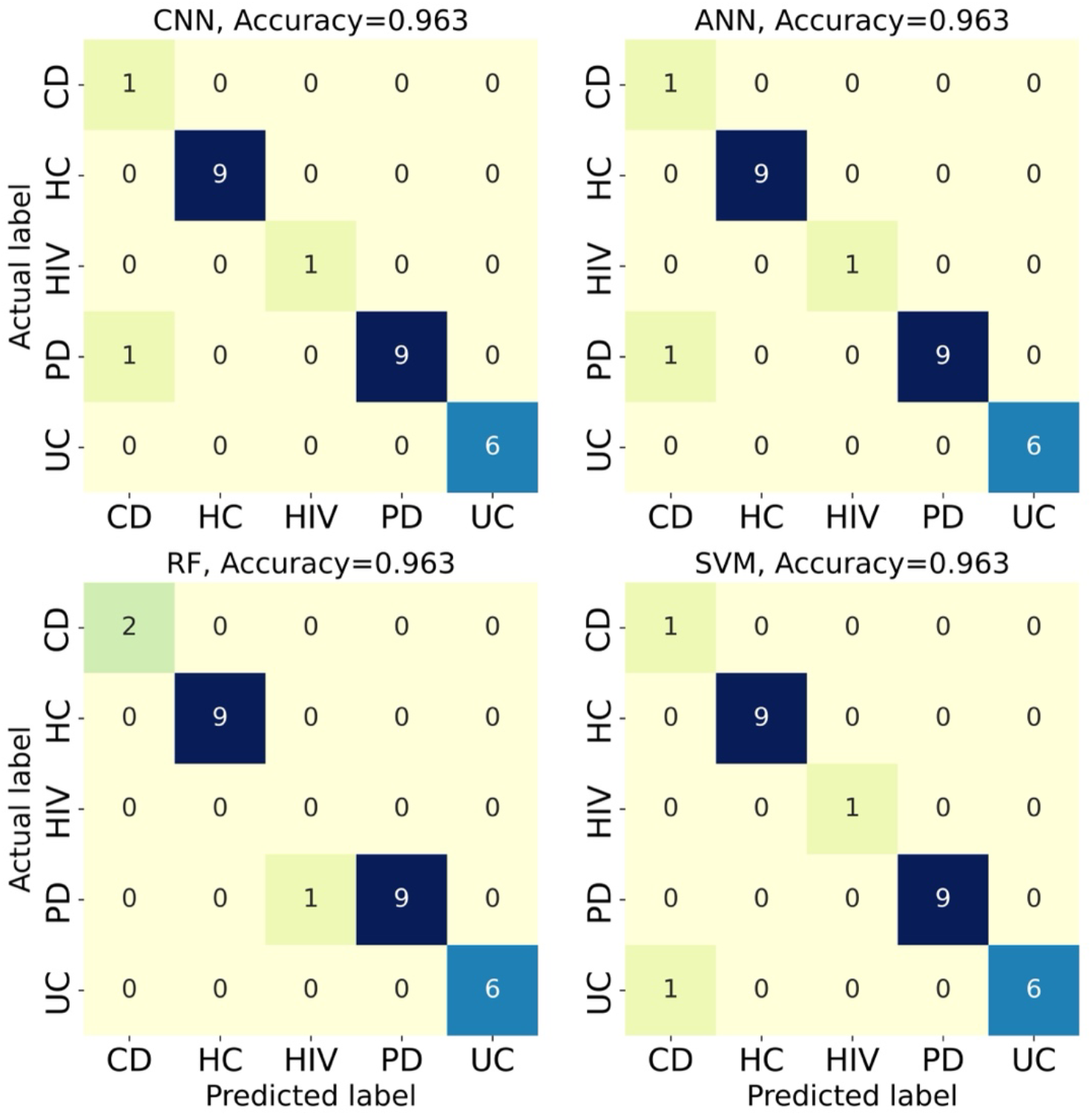
The confusion matrix for the classification of the five conditions for the baseline data

**Figure 4:**
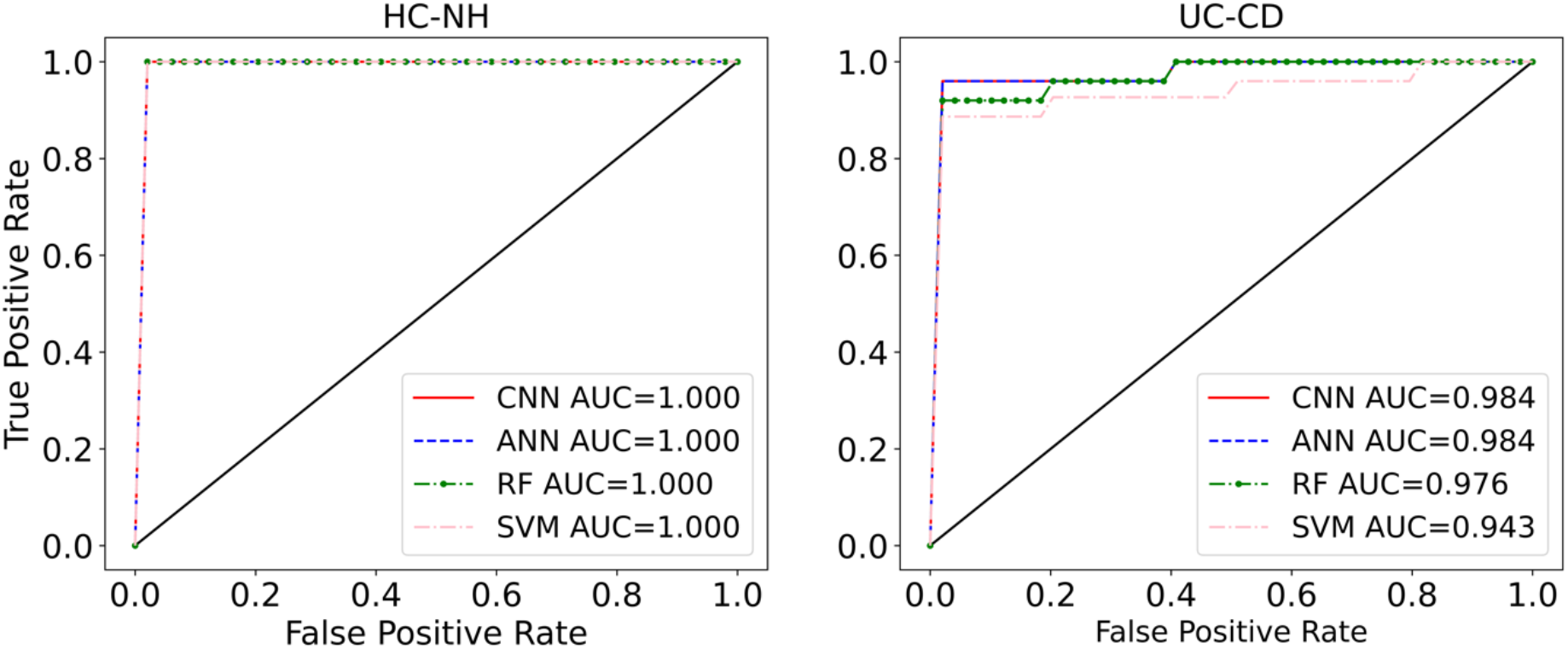
ROC curves with AUC values listed for the binary classification of HC-NH and UC-CD for the baseline data

### Prediction of the diseases based on the fiber data

Similar to the baseline data, we performed three types of classifications for the fiber data. Dietary fiber is known to modify the gut microbiota composition [34], adding another layer of intraindividual gut microbiota variability that could further differentiate classification of individuals amongst disease states. Table 2 demonstrates how the classification metrics vary for different algorithms when fiber data is introduced. We notice that RF and SVM methods lead to predictions with higher accuracy compared to ANN and CNN. Furthermore, the presence of more data points in the fiber data outweighs the microbial shifts caused by dietary fiber and significantly boosts the classification accuracy, particularly for each of the five individual conditions. As shown in Figure *5*, the heatmap for one of the folds in the 5-fold cross-validation, we note that for all methods, the only conditions that are misclassified belong to UC, CD and HIV cases. Further, we note that for the UC-CD classification, increasing the data size improved the classification accuracy compared to the baseline data. However, the algorithms still misidentify a small number of the CD cases as UC and vice versa (3 out of 88, Figure 5), which is further reflected in the receiver operating characteristic curve (ROC) curves and area under characteristic curve (AUC) values (Figure *4*), where prediction accuracy in distinguishing them reached as high as 97% (RF, Table 2).

**Table 2:** Classification performances using the fiber data

| Task | Classifier | Macro precision | Macro Recall | Macro F1 | Micro | Accuracy |
| --- | --- | --- | --- | --- | --- | --- |
| Five conditions | RF | 0.989±0.003 | 0.990±0.005 | 0.990±0.004 | 0.989±0.005 | 0.989±0.005 |
| | SVM | 0.990±0.004 | 0.985±0.007 | 0.987±0.005 | 0.988±0.005 | 0.988±0.005 |
| | ANN | 0.934±0.037 | 0.947±0.028 | 0.934±0.028 | 0.958±0.018 | 0.958±0.018 |
| | CNN | 0.981±0.005 | 0.959±0.029 | 0.968±0.018 | 0.979±0.007 | 0.979±0.007 |
| HC-NH | RF | 1.000±0.000 | 1.000±0.000 | 1.000±0.000 | 1.000±0.000 | 1.000±0.000 |
| | SVM | 0.999±0.001 | 0.998±0.004 | 0.999±0.003 | 0.999±0.002 | 0.999±0.002 |
| | ANN | 0.961±0.011 | 0.967±0.014 | 0.963±0.008 | 0.971±0.005 | 0.971±0.005 |
| | CNN | 0.992±0.006 | 0.983±0.017 | 0.987±0.012 | 0.990±0.010 | 0.990±0.010 |
| UC-CD | RF | 0.973±0.019 | 0.971±0.019 | 0.972±0.019 | 0.972±0.019 | 0.972±0.019 |
| | SVM | 0.963±0.015 | 0.963±0.015 | 0.963±0.015 | 0.963±0.015 | 0.963±0.015 |
| | ANN | 0.950±0.023 | 0.950±0.022 | 0.950±0.023 | 0.950±0.022 | 0.950±0.022 |
| | CNN | 0.963±0.018 | 0.960±0.018 | 0.961±0.018 | 0.961±0.018 | 0.961±0.018 |

**Figure 5:**
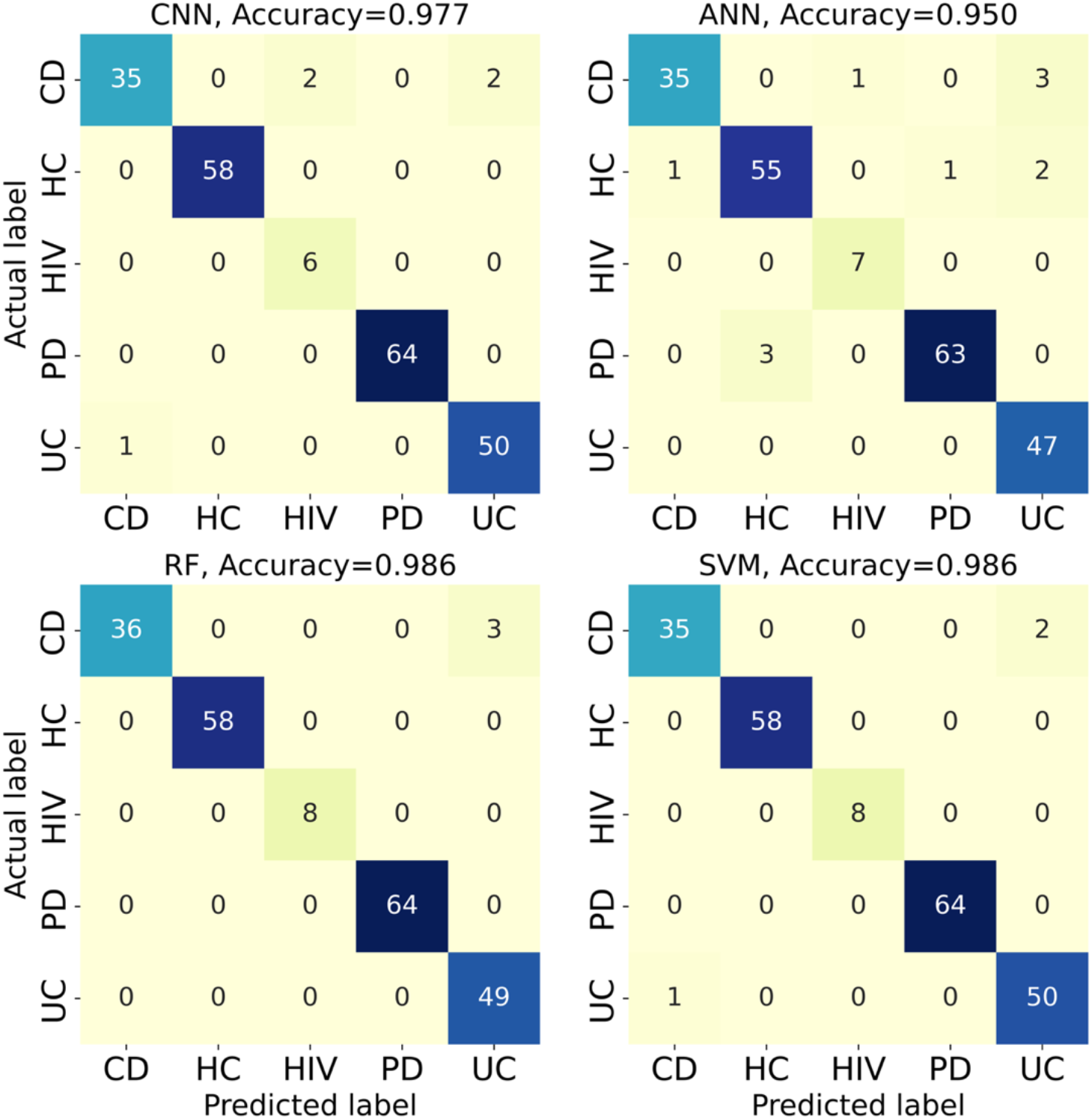
The confusion matrix for the classification of the five conditions for the fiber data

We also note that in the fiber dataset, similar to the baseline data, there is still misclassifications regarding the HIV cases, due to their low abundance in the training datasets. However, the misclassification of the PD cases is removed by increasing the data size compared to the baseline data. The highest classification accuracy belongs to the HC-NH case, suggesting that algorithms can perfectly discriminate between HC and NH even in the presence of environmental shifts that affect the gut microbial community. The ROC curves and AUC values further confirm the capability of the current ML methods, as shown in Figure *6*. As evident, the algorithms almost behave as perfect classifiers when it comes to detecting HC vs NH, which is very promising and suggests that all the four diseases used in this study induce conformational changes in the microbiome, which are conserved and can be detected by ML algorithms even in the presence of fiber treatment.

**Figure 6:**
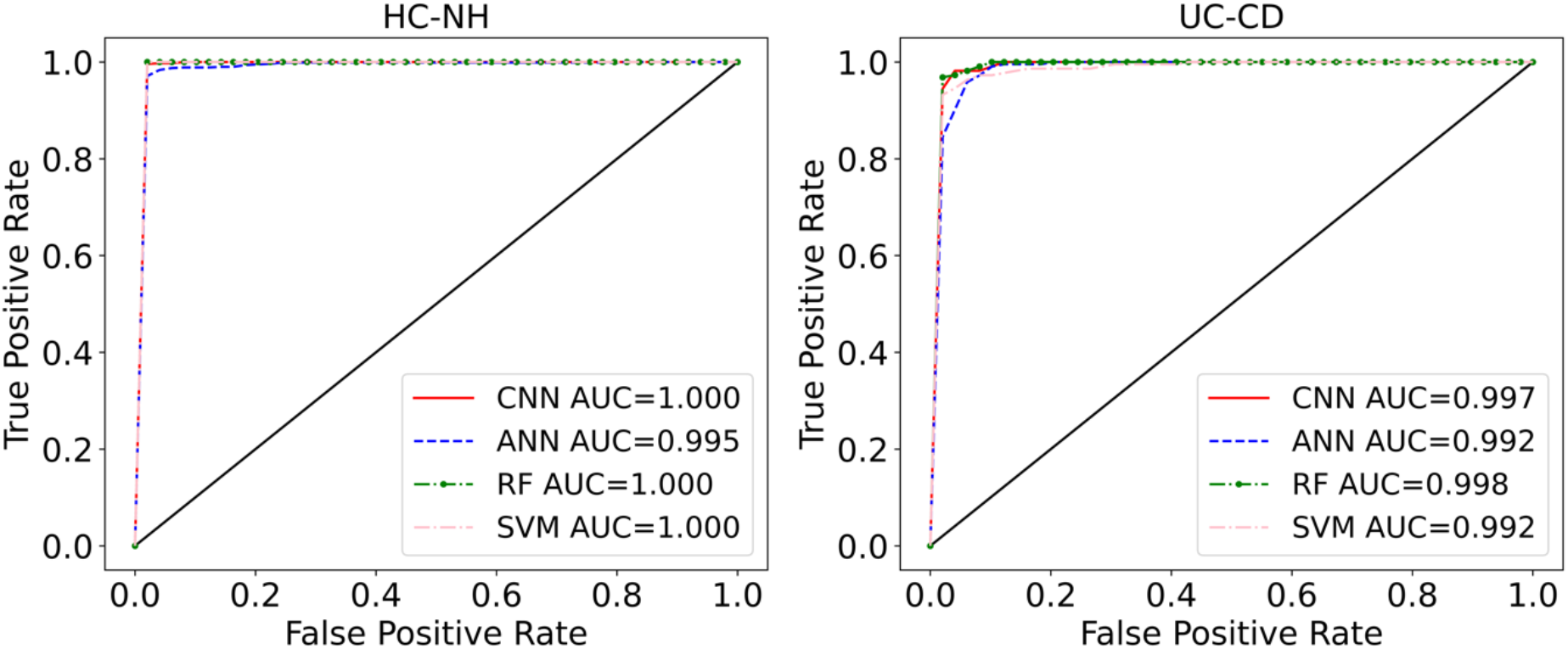
ROC curves with AUC values listed for the binary classification of HC-NH and UC-CD for the fiber data

### Visualization

To further demonstrate the strength of the ML models, we use the values of the last activation layer of ANN and conduct the principal component analysis (PCA) to visualize the data, as shown in Figure *7*. Three principal components, as opposed to two, are used to create 3-dimensional plots as we note that the third principal component, for instance, in the case of five conditions, plays an important role in visually separating the data. We note that the good distinction of different labels for both the baseline and fiber datasets is in line with the accuracy values we obtained in Table *1* and Table *2*. For instance, in the case of NH-HC, for both the baseline and fiber data, we notice there is a clear distinction between the two cases, which is in agreement with the high AUC values for both cases. These distinct separations suggest that the gut microbiota of healthy individuals and individuals in different disease states can be discriminated through machine learning even after environmental shifts (fiber fermentation), and thus could be an important tool for diagnosis of diseases that affect the gut locally and other body systems.

**Figure 7:**
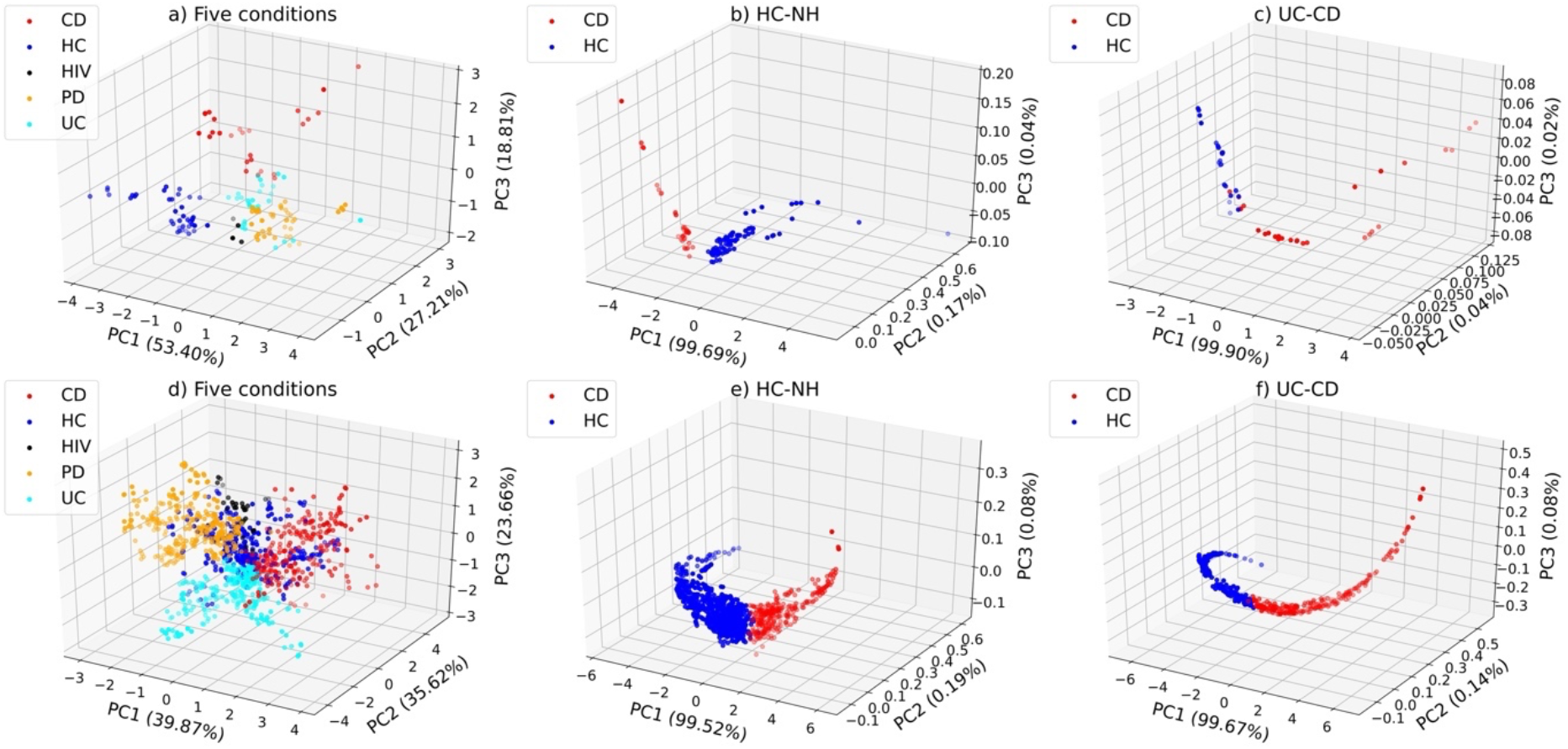
Visualization of the a) five conditions b) HC-NH c) UC-CD for the baseline data, and d) five conditions e) HC-NH f) UC-CD for the fiber data. We have used the data in the last layer of ANN to visualize the data using PCA.

## Conclusion

Studies of gut microbiota signatures often target one specific disease or state, but comparisons of the microbiota across different diseases can be challenging since some pathways that affect the gut are shared across distinct diseases. For instance, intestinal inflammation – a strong modifier of the gut microbial community [35] is a common finding in several diseases including UC, CD, PD and HIV enteropathy [36]–[38]. Moreover, there are a number of individualized physiological symptoms related to the gut microbiota within diseases that further make such classification difficult. In the current study, different ML algorithms have been applied for the classification of five disease conditions, including PD, UC, CD, HIV, and HC, in three types of classification problems. All the genomic data were preprocessed and transformed to OTU forms and then normalized. We have adopted two different types of datasets; the baseline data, where no treatments are given to patients and the fiber data, which includes both the baseline data and the data where fiber treatments are given to subjects. We have shown that it is possible to distinguish between the five conditions with accuracies as high as 95%, with SVM demonstrating the best performance among all methods. We have demonstrated that classifiers behave almost as perfect classifiers when it comes to classifying the HC and NH for both the baseline and fiber datasets. Additionally, we have shown that UC can be distinguished from CD with high accuracy using the OTU-based genomic data, even in the presence of the fiber treatment. The ability of our method to distinguish UC from CD with high accuracy has major clinical impact because current methods including invasive diagnostic tools like endoscopic procedures can fail to distinguish up to 20% of CD from UC [39]. Accurate diagnosis of CD vs UC is essential for appropriate selection of surgical approach for treatment of these patients when they require surgery. Future prospective large cohort studies are required to confirm our proposed method. As for PD, current diagnosis is primarily based on clinical symptoms and sign with associated inaccuracy of subjective diagnostic tools [40]. Brain imaging diagnostic tools are expensive, not wildly available and sensitivity/specificity are not optimal [41], [42]. Thus, if future cohort study with large number of PD patients as well as those with Parkinsonism confirm our finding, then use of non-invasive stool microbiota can be used as an objective means to accurately diagnose PD, the second most common neurodegenerative disease with alarming increase in its incidence in western societies [43]. It should be noted that our study is a proof-of-concept study and future studies with much larger sample size and use of diverse study cohort regarding age, gender, race/ethnicity and dietary habits is needed to confirm our results and the generality of our model for the diagnosis/ disease course prediction of these disorders that are associated with gut microbiota dysbiosis.

## Ethics approval and Consent to participate

All methods were carried out in accordance with relevant guidelines and regulations. All participants signed the Rush University Medical Center (RUMC) Institutional Review Board approved informed consent forms (ORA#: 07100403; 12020204; 07092603; L04092807)

## Consent for publication

Not applicable.

## Availability of data and materials

All the data and python notebook files used in this study are available in the GitHub repository https://github.com/ArezooArdekani/Classifying_diseases_gut_microbiota_ML.

## Authors’ contributions

MB conducted the machine learning analysis; TJ, MB, TC and BZ conducted data preprocessing; AK and AL collected the data; MB, TJ, AK, BH and AA wrote the manuscript; AK, BH, AA helped design the study; AL critically reviewed the manuscript; BH and AA supervised the project; BH and AA provided the funding. All authors reviewed and agreed upon the final manuscript.

## Acknowledgement

We thank Dr. Kathleen Shannon for her prior diagnosis and recruitment of Parkinson’s disease subjects at Rush University Medical Center. We thank Phillip Engen for his assistance with sample collection, preparation, and clinical data for research subjects examined.

